# *Phosphate transporter* genes: genome-wide identification and characterization in *Camelina sativa*

**DOI:** 10.1101/2021.02.28.433256

**Authors:** Soosan Hasanzadeh, Sahar Faraji, Abdullah, Parviz Heidari

## Abstract

Phosphorus is known as a key element associated with growth, energy, and cell signaling. In plants, phosphate transporters (PHTs) are responsible for moving and distributing phosphorus in cells and organs. *PHT* genes have been genome-wide identified and characterized in various plant species, however, these genes have not been widely identified based on available genomic data in *Camellia sativa*, which is an important oil seed plant. In the present study, we found 66 *PHT* genes involved in phosphate transporter/translocate in *C. sativa*. The recognized genes belonged to *PHTs1, PHTs2, PHTs4, PHOs1, PHO1 homologs, glycerol-3-PHTs, sodium dependent PHTs, inorganic PHTs, xylulose 5-PHTs, glucose-6-phosphate translocators*, and *phosphoenolpyruvate translocators*. Our finding revealed that PHT proteins are divers based on their physicochemical properties such as Isoelectric point (pI), molecular weight, GRAVY value, and exon-intron number(s). Besides, the expression profile of *PHT* genes in *C. sativa* based on RNA-seq data indicate that *PHTs* are involved in response to abiotic stresses such as cold, drought, salt, and cadmium. The tissue specific expression high expression of *PHO1* genes in root tissues of *C. sativa*. In additions, four *PHTs*, including a *PHT4;5* gene, a *sodium dependent PHT* gene, and two *PHO1 homolog 3* genes were found with an upregulation in response to aforementioned studied stresses. In the current study, we found that PHO1 proteins and their homologs have high potential to post-translation modifications such as N-glycosylation and phosphorylation. Besides, different cis-acting elements associated with response to stress and phytohormone were found in the promoter region of *PHT* genes. Overall, our results show that *PHT* genes play various functions in C. *Sativa* and regulate *Camellia* responses to external and intracellular stimuli. The results can be used in future studies related to the functional genomics of *C. sativa.*

## Introduction

Phosphorus (P) is one of the main macronutrients, which form many fundamental structures and regulate various processes of the plant cells. It serves the cell as a constitutive part of DNA, RNA, and membrane phospholipids, as a key element in energy metabolism and transfer, and as a component in signal transduction, photosynthesis, the metabolism of sugars, and respiration (Raghothama, 2000). Plants uptake only the inorganic phosphorus forms (Pi: H_2_PO_4_^-^ and HPO_4_^2-^). The slow rate of diffusion and high interaction of P with other elements (like Al, Fe, and Ca) in the soil make it least available macronutrient in the rhizosphere (Smith et al., 2003). The concentration of available P in soil barely gets to 10 μM, while concentration of Pi in plant cells is generally more than 10 mM (Tran et al., 2010). Therefore one of the most specialized plant strategies to acquire Pi is phosphate transporters (PHTs) (Smith, 2002). Two systems for Pi transporters in plants have been identified: The high-affinity phosphate transporters function at μM range of Pi concentration which is the usual Pi concentration in cultivated soils (Poirier and Bucher, 2002). On the other hand, the low-affinity phosphate transporters operate at mM range of Pi concentration (Karthikeyan et al., 2002). Plants employ multiple PHT families including PHT1, PHT2, PHT3, PHT4, PHT5, and PHO1 (Yang et al., 2020; Zhang et al., 2016). PHT1 is a Pi:H^+^ symporter with a high affinity for Pi, which is localized in the plasma membrane and belongs to the major facilitator superfamily (MFS) (Nussaume et al., 2011). *AtPHT1* was the first *PHTs* identified in higher plants (Muchhal et al., 1996). In *Arabidopsis thaliana,* the *PHT1* family consists of nine homologs which eight of them are expressed in root tissues (Młodzińska and Zboińska, 2016). Among PHT1 homologs, AtPHT1;1 and AtPHT1;4 play significant roles in Pi uptake under both sufficient and deficient pi concentrations (Shin et al., 2004). AtPHT1;8 and AtPHT1;9 participate in Pi translocation from root to shoot and both function in phosphate uptake under Pi deficiency (Remy et al., 2012). In *Oryza sativa,* 13 homologs for *OsPHT1* have been identified which all of them are expressed in rice roots (Jia et al., 2011). A recent study by upregulation of seven *PHT1* subfamily genes in rice showed that phosphate uptake from the soil under phosphorus deficiency considerably relies on *PHT1* (Gho and Jung, 2019). PHT2 is a low-affinity Pi:H^+^ symporter present in the chloroplast envelope (Daram et al., 1999). The activity of PHT2 impacts on Pi allocation inside the plant; it also moderates the plant’s responses to Pi starvation through translocating of Pi inside the leaves and induction and expression of Pi-starvation response genes (Versaw and Harrison, 2002). PHT3 or Mitochondrial Phosphate Transporter (MPT) family is a high-affinity PHT, which is localized to the inner mitochondrial membrane and Its function was reported to be Pi:H^+^ symporter and Pi:OH^-^ antiporter (Lee and Yoon, 2018). It is PHT3 responsibility to provide the required phosphate for ATP synthesis by transporting Pi from the cytosol into the mitochondrial matrix (Yu et al., 2018). Three homologs for Arabidopsis *PHT3 (AtPHT3;1-3)* and six for rice *PHT3* genes have been identified (Huang et al., 2020). The expression of *PHT4* genes takes place both in roots and leaves. PHT4;1 is a thylakoid membrane PHT, which appears to be involved in defense against pathogens (Guo et al., 2008), while PHT4;2 transports Pi within root’s heterotrophic plastids and affects starch accumulation as well (Sun et al., 2017). PHT4;4 is an ascorbate transporter located in the chloroplast inner envelope membrane, which is associated with protection from high light stress (Miyaji et al., 2015). PHT4;6 is localized to the Golgi apparatus and cooperates with salt tolerance (Cubero et al., 2009). *Oryza sativa* also has six *PHT4* genes, which are very well described in a previous study (Ruili et al., 2020). PHT5 family acts as a vacuolar PHT and consists of three members in Arabidopsis (Liu et al., 2016). Regarding PHO1, using a map-based positional cloning strategy led to the identification of this gene family in Arabidopsis (Hamburger et al., 2002). The Arabidopsis genome contains 11 members of the *PHO1* gene family, which are localized to the endomembrane (majorly Golgi) of root pericyclic cells (Gu et al., 2016; Liu et al., 2012). The expression pattern of all *PHO1 homologs* proposes that *PHO1,* in addition to transporting Pi to the vascular cylinders in various tissue, also has a probable role in phosphate acquiring into cells, like pollen or root epidermal/cortical cells (Wang et al., 2004). Under Pi deficiency, *PHO1* also plays a part in the long distance (root-to-shoot) signal transduction cascade (Młodzińska and Zboińska, 2016).

*Camelina sativa* is an oilseed member of Brassicaceae, which has gained great attention due to its high potential for biofuel production (Ahmadizadeh et al., 2020b; Murphy, 2016). Also, *C. sativa* is resistant to doughtiness, salinity and coldness and many pathogens (Brock et al., 2018). A study on *C. sativa* seed yield response to various fertilizers showed that maximum predicted seed yield was obtained without using Pi fertilizer (Solis et al., 2013). To reduce reliance on Pi fertilizers and ensuring sustainable agriculture, it is required to identify and characterize PHTs especially in low input crops such as *C. sativa*, to be able to develop Pi-efficient crops. Genomic analyses of PHT families have been conducted on various plants, including Arabidopsis, rice (Liu et al., 2011), poplar (Zhang et al., 2016), Apple (Sun et al., 2017), sorghum (Wang et al., 2019), and rapeseed (Yang et al., 2020). However, none of the report available for C. *sativa.* In this study, we conduct a comprehensive genome-wide analysis to identify and characterize *PHT* gene families in *C. sativa* and provide functional dissection of *CsPHTs.*

## Materials and methods

### Recognition of phosphate transporter family members in *Camelina sativa*

PHT proteins of the model plant Arabidopsis were used as queries in BLASTP tool search (E-value <1e-5) from the Ensembl Plants database (Bolser et al., 2017) to identify and retrieve genes encoded phosphate transporter (PHT) proteins in *C. sativa.* The identified PHTs were analyzed using Pfam (Finn et al., 2010) and Bach CDD-search of NCBI database (Marchler-Bauer et al., 2015) to check the presence of the common conserved domains and avoid truncated protein. We also retrieved sequences of genomic DNA, cDNA, and promoter regions. Furthermore, the ProtParam tool of ExPASy (Gasteiger et al., 2005) was applied to predict the isoelectric points (*pI*), molecular weights (MW), and the grand average of hydropathy (GRAVY).

### Evolutionary analysis

For the evolutionary study, the protein sequences of *phosphate transporter* genes in *C. sativa* along with their orthologues in model plants, rice and Arabidopsis, were aligned by multiple alignment using the clustalW method. The phylogenetic relationship of the phosphate transporter genes was created using the unrooted neighbor-joining method of MEGA X (Kumar et al., 2018) with 1000 bootstrap replicates and finally, the created phylogeny tree was improved using an interactive tree of life (Letunic and Bork, 2019).

### Gene expression

To study the expression profile of *phosphate transporter* genes, the available RNA-seq data related to *C. sativa* was screened to determine the expression levels of the *phosphate transporter* family members. In the present study, the RNA-seq data in leaf, flower, stem, and root of *C. sativa* were selected from the NCBI database with available accessions SRR935362, SRR935369, SRR935365, and SRR935368, respectively. Besides, RNA-seq data related to abiotic stresses, including drought stress (SRR935380), NaCl stress (SRR935382), cold stress (SRR935372), cadmium stress (SRR935383), and normal condition (SRR935385), were evaluated to state the expression pattern of *phosphate transporter* genes in *C. sativa*. The transcript levels of *phosphate transporter* genes were calculated using FPKM (fragments per kilobase of exon model per million mapped reads). Finally, heatmaps were created using log2 transformed by TBtools software (Chen et al., 2020).

### Promoter region analyses and prediction of post-translational modifications of PHT proteins

The promoter region (1500 bps upstream of the start codon) of *phosphate transporter* genes in *C. sativa* was screened to find key cis-regulatory elements related to response to stresses and hormones using the PlantCare database (Lescot et al., 2002). The phosphorylation and N-glycosylation modification as important types of post-translational modifications were predicted in phosphate transporter proteins in *C. sativa.* The potential phosphorylation site was estimated using NetPhos 3.1 server (Blom et al., 2004) with potential value > 0.5 and NetNGlyc 1.0 server (Gupta and Brunak, 2002) was used to predict the N-glycosylation sites into the amino acid sequence of phosphate transporter proteins.

## Results

In the genome of *C. sativa*, we identified sixty six non redundant *PHT* genes, including eight *PHT1* genes, three *PHT2* genes, ten *PHT4* genes, three *PHO1* genes, nineteen *PHO1 homolog* genes, eight *glycerol-3-PHT* genes, three *sodium dependent PHT* genes, four *probable inorganic PHT* genes, three *xylulose 5-PHT* genes, three *glucose-6-phosphate translocator* genes, and two *phosphoenolpyruvate translocator* genes (Table 1). The recognized PHT proteins showed a high diversity based on physicochemical properties such as pI, molecular weight, GRAVY value, exon number, and protein length. Exon number of studied PHTs varied from 1 to 20 and protein length ranged from 209 to 1230 aa. Furthermore, pI showed a variation between 5.11 and 10.11 in phosphate transporter/translocate proteins of *C. sativa.* Besides, the GRAVY value revealed that all PHO homolog proteins are hydrophilic proteins. The studied PHT genes were distributed on 20 chromosomes of *C. sativa* (Fig. 1). Most phosphate transporter genes were located in chromosome 6 (eight genes) and chromosome 9 (nine genes). Chromosomes 1, 15, and 19 each had a *PHO* gene, and chromosomes 2 and 12 had a *glycerol-3-PHT* gene. Besides, the location (start and end position) of each *phosphate transporter/translocate* gene on the Camelina chromosome is also mentioned in Table 1. Our results revealed the unequal distribution of *phosphate transporter/translocate* genes within *C. sativa* genome.

**Table 1.** Properties of recognized *phosphate transporter* genes into *C. sativa* genome

| Gene ID | Description | Chr. location | CDS (bp) | Exon | Peptide (aa) | MW (kDa) | pI | GRAVY |
| --- | --- | --- | --- | --- | --- | --- | --- | --- |
| *Csa05g014350* | PHT1 | 5:5014921-5018711 | 2223 | 2 | 534 | 58.73 | 8.35 | 0.294 |
| *Csa04g052890* | PHT1 | 4:25092914-25096711 | 1381 | 2 | 392 | 42.75 | 7.00 | 0.455 |
| *Csa06g042880* | PHT1 | 6:21171097-21174709 | 2275 | 2 | 534 | 58.69 | 8.35 | 0.299 |
| *Csa09g081170* | Putative PHT | 9:30797478-30800557 | 1593 | 2 | 530 | 58.51 | 8.54 | 0.466 |
| *Csa07g047990* | Putative PHT | 7:24682311-24685313 | 1590 | 2 | 529 | 58.36 | 8.93 | 0.454 |
| *Csa16g040570* | Putative PHT | 16:20731839-20736565 | 3305 | 2 | 530 | 58.47 | 8.54 | 0.478 |
| *Csa03g024720* | PHT1;8 | 3:9470369-9472993 | 1611 | 2 | 536 | 59.10 | 6.52 | 0.381 |
| *Csa14g026060* | PHT1;8 | 14:10079381-10081989 | 1611 | 2 | 536 | 59.08 | 6.46 | 0.385 |
| *Csa06g006270* | PHT2;1 | 6:3480684-3482786 | 1880 | 3 | 587 | 61.22 | 9.34 | 0.554 |
| *Csa09g010040* | PHT2;1 | 9:4351633-4354047 | 2201 | 3 | 587 | 61.27 | 9.34 | 0.558 |
| *Csa04g012130* | PHT2;1 | 4:5383653-5386072 | 2192 | 3 | 587 | 61.32 | 9.39 | 0.534 |
| *Csa06g040800* | PHT4;2 | 6:20643394-20647032 | 1892 | 8 | 509 | 55.59 | 9.68 | 0.515 |
| *Csa05g015490* | PHT4;2 | 5:5626753-5628888 | 1486 | 7 | 393 | 42.82 | 8.94 | 0.703 |
| *Csa04g051740* | PHT4;2 | 4:24495983-24499701 | 1948 | 8 | 509 | 54.97 | 9.74 | 0.547 |
| *Csa13g023330* | PHT4;5 | 13:9052618-9057824 | 1482 | 16 | 209 | 22.98 | 8.46 | 0.001 |
| *Csa20g032330* | PHT4;5 | 20:10882393-10888480 | 1503 | 16 | 500 | 54.28 | 5.11 | 0.349 |
| *Csa08g014220* | PHT4;5 | 8:5926491-5931307 | 1882 | 15 | 525 | 57.55 | 6.43 | 0.252 |
| *Csa09g045340* | PHT4;3 | 9:16771385-16774489 | 2138 | 9 | 533 | 58.18 | 9.66 | 0.464 |
| *Csa06g020780* | PHT4;3 | 6:11456825-11460071 | 2112 | 9 | 655 | 71.02 | 9.68 | 0.468 |
| *Csa11g070280* | PHT4 | 11:33280938-33283592 | 2041 | 2 | 432 | 47.08 | 9.44 | 0.678 |
| *Csa20g071480* | PHT4 | 20:24416985-24419803 | 2110 | 2 | 432 | 47.10 | 9.44 | 0.673 |
| *Csa09g096350* | Glucose-6-phosphate translocator2 | 9:36509054-36511464 | 1661 | 5 | 388 | 42.71 | 9.70 | 0.396 |
| *Csa07g063090* | Glucose-6-phosphate translocator2 | 7:31985709-31987975 | 1586 | 5 | 389 | 42.81 | 9.66 | 0.387 |
| *Csa16g053570* | Glucose-6-phosphate translocator2 | 16:27479855-27482074 | 1491 | 5 | 403 | 44.50 | 9.61 | 0.347 |
| *Csa13g020410* | Xylulose 5- PHT | 13:7578890-7580469 | 1580 | 1 | 414 | 45.43 | 9.74 | 0.335 |
| *Csa20g025800* | Xylulose 5- PHT | 20:8621980-8623242 | 1263 | 1 | 420 | 46.16 | 9.74 | 0.285 |
| *Csa08g009840* | Xylulose 5- PHT | 8:4008763-4010025 | 1263 | 1 | 420 | 46.17 | 9.78 | 0.309 |
| *Csa09g064460* | Probable inorganic PHT1-7 | 9:24544974-24548295 | 1608 | 2 | 535 | 58.63 | 9.05 | 0.457 |
| *Csa04g040820* | Probable inorganic PHT1-7 | 4:19796739-19799556 | 1938 | 2 | 535 | 58.37 | 8.68 | 0.325 |
| *Csa06g029160* | Probable inorganic PHT1-7 | 6:16352706-16355582 | 1981 | 2 | 535 | 58.42 | 8.68 | 0.325 |
| *Csa18g010490* | Probable inorganic PHT1-6 | 18:5437712-5444775 | 4707 | 6 | 998 | 109.04 | 9.20 | 0.455 |
| *Csa09g047740* | Glycerol-3- PHT1 | 9:17201598-17204706 | 2225 | 3 | 526 | 56.45 | 6.32 | 0.411 |
| *Csa06g021070* | Glycerol-3- PHT1 | 6:11705560-11708615 | 2031 | 3 | 526 | 56.44 | 6.32 | 0.424 |
| *Csa11g019360* | Glycerol-3- PHT2 | 11:8685846-8687786 | 1857 | 2 | 501 | 54.33 | 6.66 | 0.582 |
| *Csa10g017760* | Glycerol-3- PHT2 | 10:7441422-7443017 | 1512 | 2 | 503 | 54.54 | 7.75 | 0.576 |
| *Csa12g028010* | Glycerol-3- PHT2 | 12:8515749-8517468 | 1636 | 2 | 502 | 54.39 | 6.66 | 0.575 |
| *Csa18g026090* | Glycerol-3-PHT1 | 18:13795519-13798239 | 1592 | 5 | 394 | 43.12 | 9.63 | 0.411 |
| *Csa02g057110* | Glycerol-3-PHT1 | 2:21239422-21241966 | 1386 | 5 | 373 | 40.91 | 9.39 | 0.360 |
| *Csa11g087830* | Glycerol-3-PHT1 | 11:42451060-42453885 | 1807 | 6 | 417 | 45.81 | 9.56 | 0.349 |
| *Csa01g028000* | PHO1 | 1:11864045-11869788 | 2343 | 12 | 653 | 74.27 | 9.74 | 0.022 |
| *Csa15g044550* | PHO1 | 15:15190897-15198630 | 3771 | 20 | 1022 | 117.10 | 9.44 | 0.131 |
| *Csa19g033930* | PHO1 | 19:13491847-13499274 | 2962 | 14 | 752 | 86.41 | 9.67 | -0.052 |
| *Csa07g036190* | PHO1 homolog10 | 7:19504364-19507950 | 2446 | 13 | 759 | 88.96 | 9.03 | -0.229 |
| *Csa05g086190* | PHO1 homolog10 | 5:31052820-31057418 | 3343 | 16 | 1055 | 122.84 | 8.00 | -0.343 |
| *Csa06g003260* | PHO1 homolog2 | 6:1774457-1778037 | 2322 | 12 | 773 | 89.73 | 9.60 | -0.120 |
| *Csa11g019220* | PHO1 homolog4 | 11:8636753-8639851 | 2238 | 11 | 745 | 87.07 | 9.34 | -0.076 |
| *Csa16g030840* | PHO1 homolog10 | 16:16337025-16340408 | 2364 | 13 | 787 | 91.35 | 8.85 | -0.217 |
| *Csa09g005530* | PHO1 homolog2 | 9:2199317-2207086 | 3732 | 20 | 1239 | 143.17 | 9.49 | -0.206 |
| *Csa17g063450* | PHO1 homolog8 | 17:21903188-21907430 | 3259 | 12 | 750 | 87.40 | 9.15 | -0.097 |
| *Csa14g017010* | PHO1 homolog3 | 14:6684564-6689160 | 2741 | 12 | 817 | 94.49 | 9.34 | -0.205 |
| *Csa03g017780* | PHO1 homolog3 | 3:6394132-6398776 | 2894 | 12 | 780 | 89.91 | 9.44 | -0.154 |
| *Csa04g018460* | PHO1 homolog9 | 4:9232367-9236955 | 2676 | 17 | 877 | 100.81 | 9.28 | -0.073 |
| *Csa06g003190* | PHO1 homolog5 | 6:1723235-1730512 | 2926 | 12 | 840 | 96.92 | 8.72 | -0.223 |
| *Csa04g008740* | PHO1 homolog5 | 4:2991897-2996176 | 3027 | 12 | 839 | 96.84 | 8.68 | -0.236 |
| *Csa09g005520* | PHO1 homolog5 | 9:2187493-2192729 | 2980 | 12 | 830 | 95.74 | 8.52 | -0.198 |
| *Csa17g019420* | PHO1 homolog3 | 17:6459607-6470223 | 2878 | 12 | 816 | 94.16 | 9.31 | -0.180 |
| *Csa03g038920* | PHO1 homolog8 | 3:17838193-17841965 | 2789 | 12 | 751 | 87.64 | 9.14 | -0.123 |
| *Csa10g017620* | PHO1 homolog4 | 10:7392473-7399374 | 3730 | 20 | 1171 | 133.75 | 7.52 | -0.460 |
| *Csa09g022610* | PHO1 homolog9 | 9:8854639-8859234 | 2412 | 13 | 739 | 85.64 | 9.39 | -0.286 |
| *Csa17g034160* | PHO1 homolog7 | 17:12430216-12433887 | 2697 | 12 | 751 | 87.70 | 9.26 | -0.051 |
| *Csa05g085290* | PHO1 homolog1 | 5:30379302-30383384 | 2700 | 12 | 784 | 90.72 | 9.00 | -0.212 |
| *Csa05g034250* | Sodium-dependent PHT1 | 5:12660817-12663456 | 1862 | 9 | 509 | 56.38 | 9.08 | 0.282 |
| *Csa07g014740* | Sodium-dependent PHT1 | 7:6596929-6599515 | 1772 | 9 | 509 | 56.23 | 9.09 | 0.297 |
| *Csa16g015150* | Sodium-dependent PHT1 | 16:5712825-5714991 | 1515 | 8 | 504 | 56.15 | 9.31 | 0.284 |
| *Csa10g038420* | Phosphoenolpyruvate translocator1 | 10:17811762-17814549 | 1810 | 9 | 449 | 48.73 | 10.08 | 0.520 |
| *Csa11g044500* | Phosphoenolpyruvate translocator1 | 11:20839051-20842672 | 1869 | 9 | 408 | 44.21 | 10.11 | 0.449 |

**Fig. 1.**
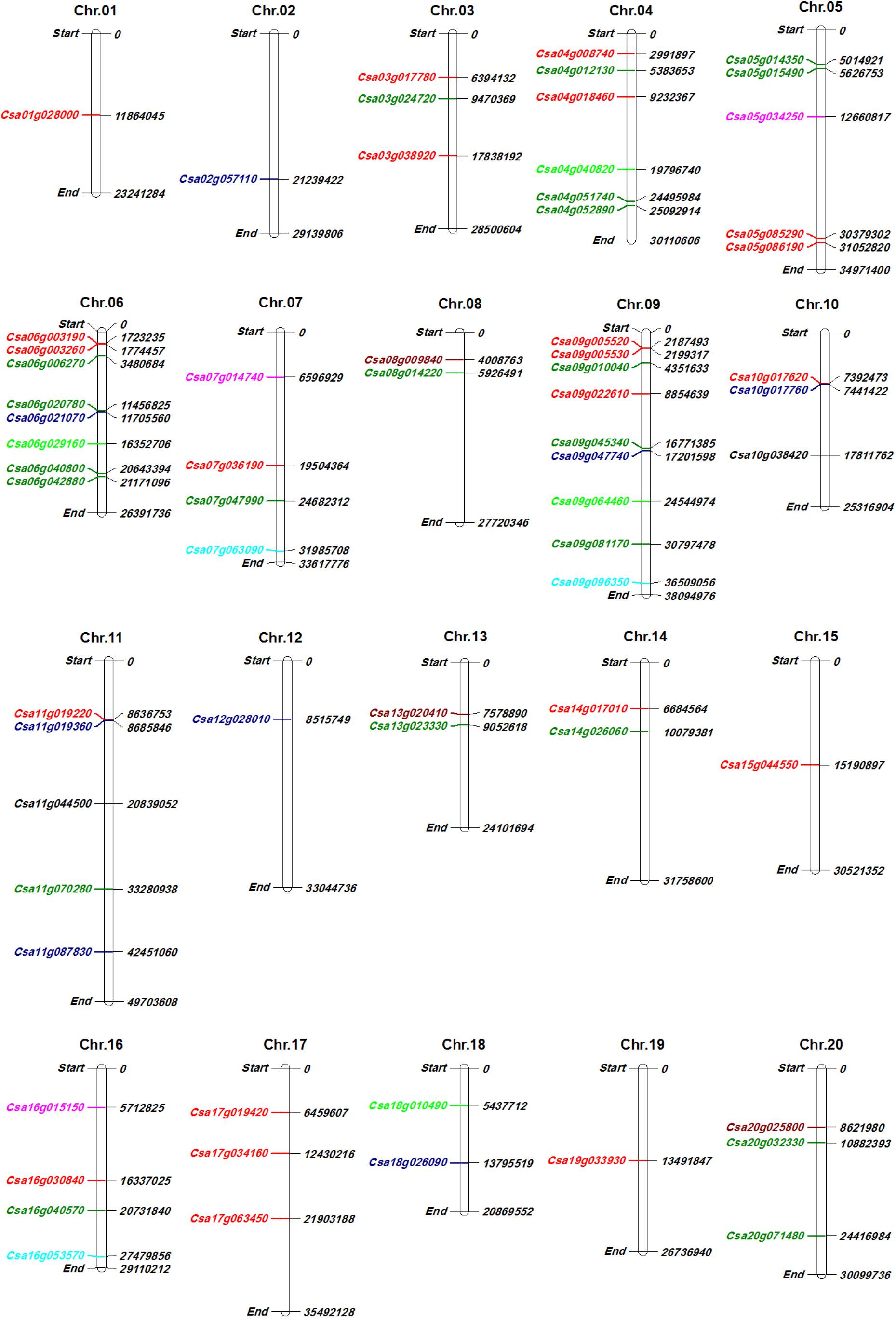
Chromosome distribution of *PHT* genes in *C. sativa* genome. Different colors used to show the type of phosphate transporter, *PHOs1* and *PHO1 homologs* (red color), *PHTs1, PHTs2,* and *PHTs4* (green color), *glycerol-3-PHTs* (blue color), *sodium dependent PHTs* (pink color), *probable inorganic PHTs* (light green), *xylulose 5-PHTs* (brown color), *glucose-6-phosphate translocators* (aqua color), and *phosphoenolpyruvate translocators* (black color).

### Phylogenetic relationships of PHTs

In the present study, a phylogenetic tree was reconstructed among phosphate *transporter/translocate* proteins of *C. sativa*,, *Arabidopsis thaliana* and *Oryza sativa* (Fig. 2). The phylogenetic analysis classified PHTs into seven different groups. Two phosphoenolpyruvate translocators, three xylulose 5-PHTs, three glucose-6-phosphate translocators as well as three glycerol-3-PHTs of *C. sativa* with their homologs of Arabidopsis and rice were located in the group I. Interestingly, all PHOs of *C. sativa* with their homologous genes were placed in the group II, while group III consisted of rice PHTs. Besides, three PHT2;1 proteins were located in group IV and three PHTs1, three putative PHTs, two PHTs1;8 with four probable inorganic PHTs were placed in group V. In addition, two glycerol-3-PHTs1 and three glycerol-3-PHTs2 of *C. sativa* with their homologs were located in the group VI. In the group VII, three sodium dependent PHTs showed more relationships with PHT4 proteins. The results revealed that PHTs of *C. sativa* are more similar to their homologs in Arabidopsis and PHTs in rice, as a monocot model plant, have high diversity than studied dicot species.

**Fig.2.**
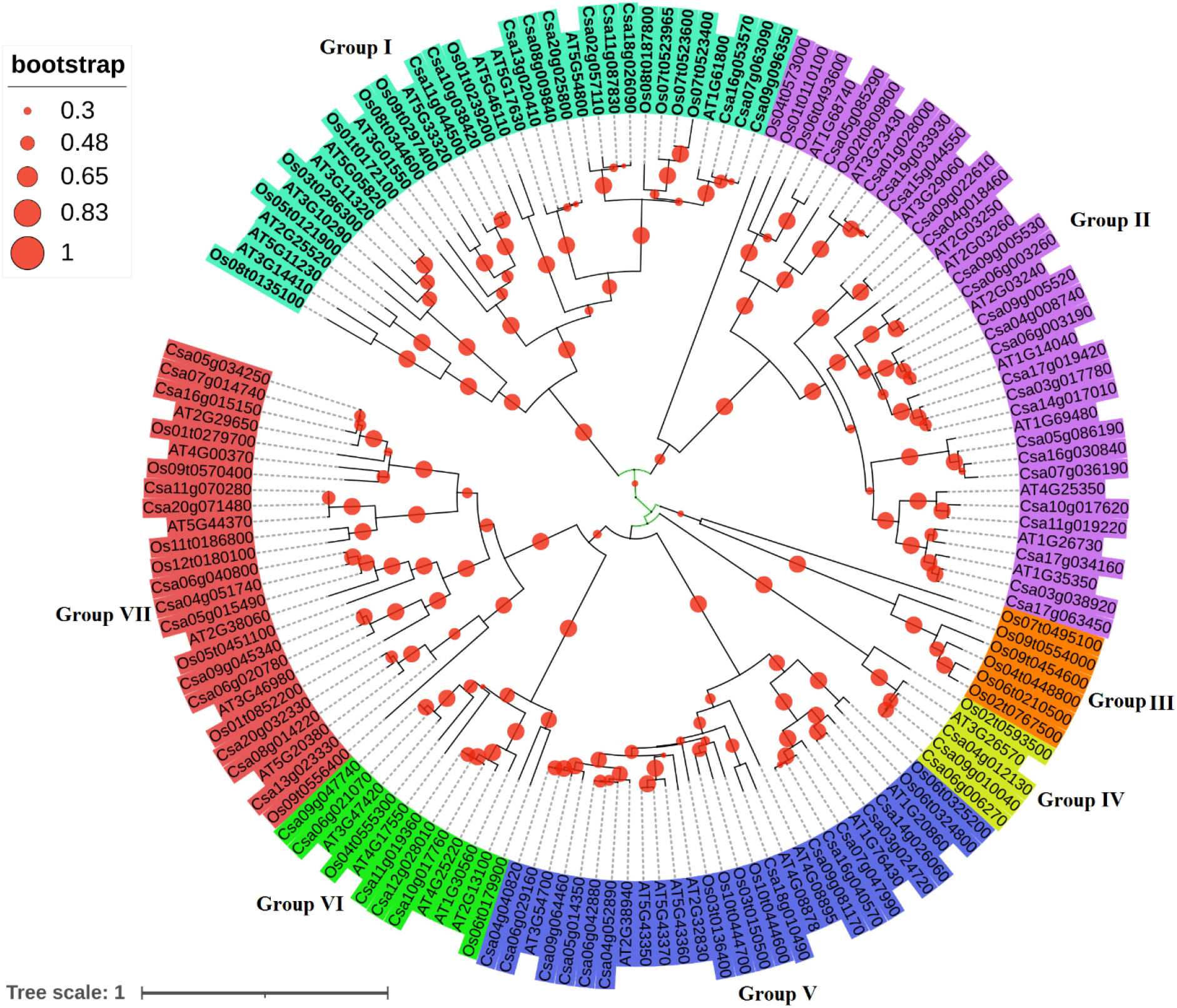
Phylogenetic tree of phosphate transporter proteins in *C. sativa* with their orthologous in Arabidopsis and rice. Phylogenetic tree was created using NJ method with 1000 bootstrap.

### Expression profile of *PHT* genes

The expression level of *PHT* genes in *C. sativa* was evaluated in different tissues and in response to abiotic stresses, including cold, NaCl, cadmium, and drought conditions (Fig. 3a, b). Regarding the expression results, *PHTs* showed diverse expression patterns in various tissues of *C. sativa* (Fig. 3a). Furthermore, our findings revealed that *PHTs* are more expressed in shoot tissues than root tissues of *C. sativa* (Fig. 4a). Besides, *PHO1* genes showed high expression in root tissues than shoot tissues. The expression profile of *PHTs* under abiotic stresses stated that *PHTs* are involved in response to drought, cold, salinity, and cadmium stress (Fig. 3b). However, *PHTs* genes showed differential expression based on the type of stress. For instance, *PHTs* were more up-regulated in response to cold stress, while they were more down-regulated in response to NaCl stress (Fig. 4b). In addition, Venn diagrams also showed differential expression patterns of *PHTs* in response to abiotic stresses (Fig. 4c, d). Regarding the Venn diagrams, four *PHTs,* including *Csa13g023330* (as a *PHT4;5* gene), *Csa05g034250* (as a *sodium dependent PHT* gene), and two *PHO1 homolog3* genes *(Csa14g017010* and *Csa17g019420)* were up regulated in response to all studied abiotic stresses (Fig. 4c). Moreover, a *PHO1 homolog5* (*Csa09g005520*) was identified as a common repressed gene in response to abiotic stresses in *C. sativa* (Fig. 4d). Overall, our findings stated that *PHTs* are involved in regulating Camelina’s responses under environmental stress conditions.

**Fig. 3.**
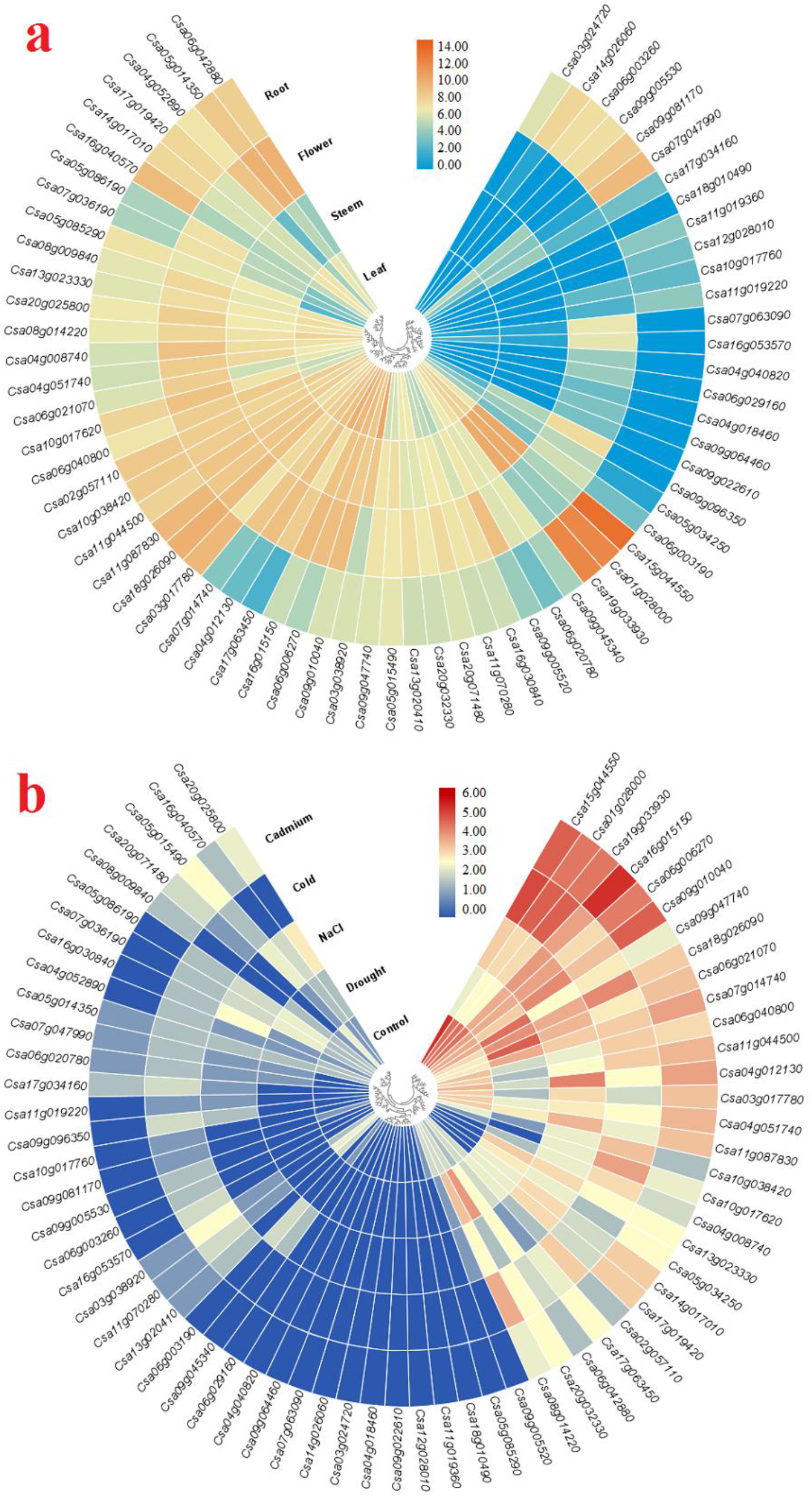
Expression profile of *phosphate transporter* genes in *C. sativa* at different tissues (a), and response to abiotic stress (b).

**Fig. 4.**
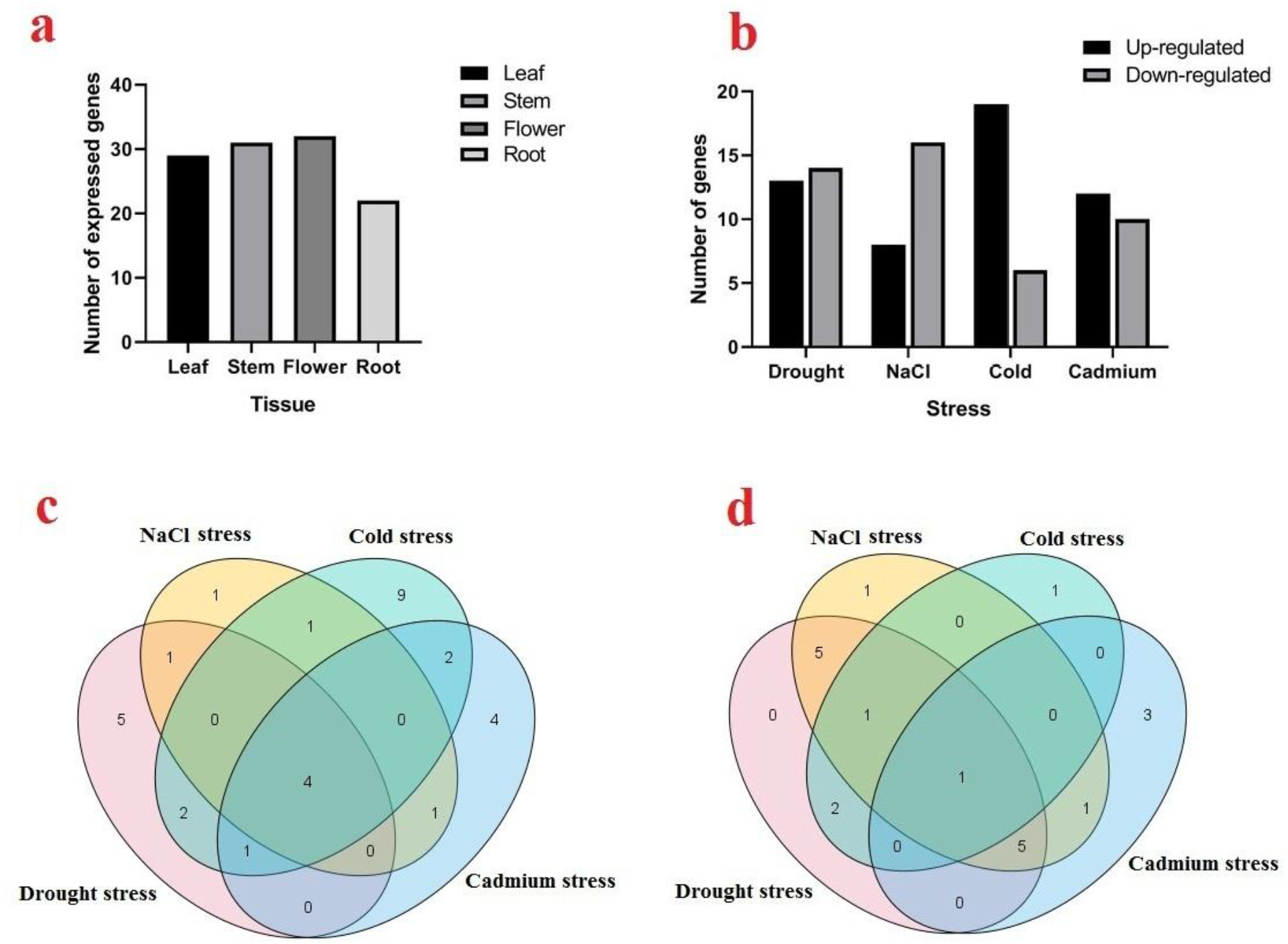
The number of expressed *PHTs* in different tissues of *C. sativa* (a), number of up-regulated and down regulated *PHTs* in *C. sativa* genomes in response to abiotic stresses, including drought, NaCl, cold, and cadmium (b), Venn diagram of up-regulated *PHT* genes (c), and down-regulated *PHT* genes in response to abiotic stresses (d).

### Post-translational modifications

In the current study, we predicted the potential site of post-translation modifications in terms of phosphorylation and N-glycosylation modifications into PTH proteins of *C. sativa* (Fig. 5). Regarding the prediction N-glycosylation results, seven PHTs, including two PHT2;1 proteins (Csa06g006270 and Csa09g010040), two PHT4;3 proteins (Csa09g045340 and Csa06g020780), two PHT4 proteins (Csa11g070280 and Csa20g071480), and a PHO1 homolog10 protein (Csa16g030840), were found non site for N-glycosylation (Fig. 5a). Besides, a PHO1 homolog10, Csa03g017780, with seven predicted sites was found as a hyper glycosylation protein and six N-glycosylation sites were predicted in four PHO1 proteins, including PHO1 homolog4 (Csa10g017620), PHO1 homolog2 (Csa09g005530), PHO1 homolog7 (Csa17g034160), and PHO1 homolog3 (Csa14g017010). In addition, PHT proteins in *C. sativa* varied from 20 to 112 sites based on potential phosphorylation modifications (Fig. 5b). A PHT4;5 protein, Csa13g023330, showed a minimum potential phosphorylation site (20 predicted sites), while two PHO1 homolog proteins, including PHO1 homolog4 (Csa10g017620), PHO1 homolog2 (Csa09g005530), were found with high potential sites (112 sites). Overall, the results of prediction of post-translation modifications revealed that PHO1 homolog proteins have more potential for phosphorylation and glycosylation than other PHT proteins in *C. sativa*.

**Fig. 5.**
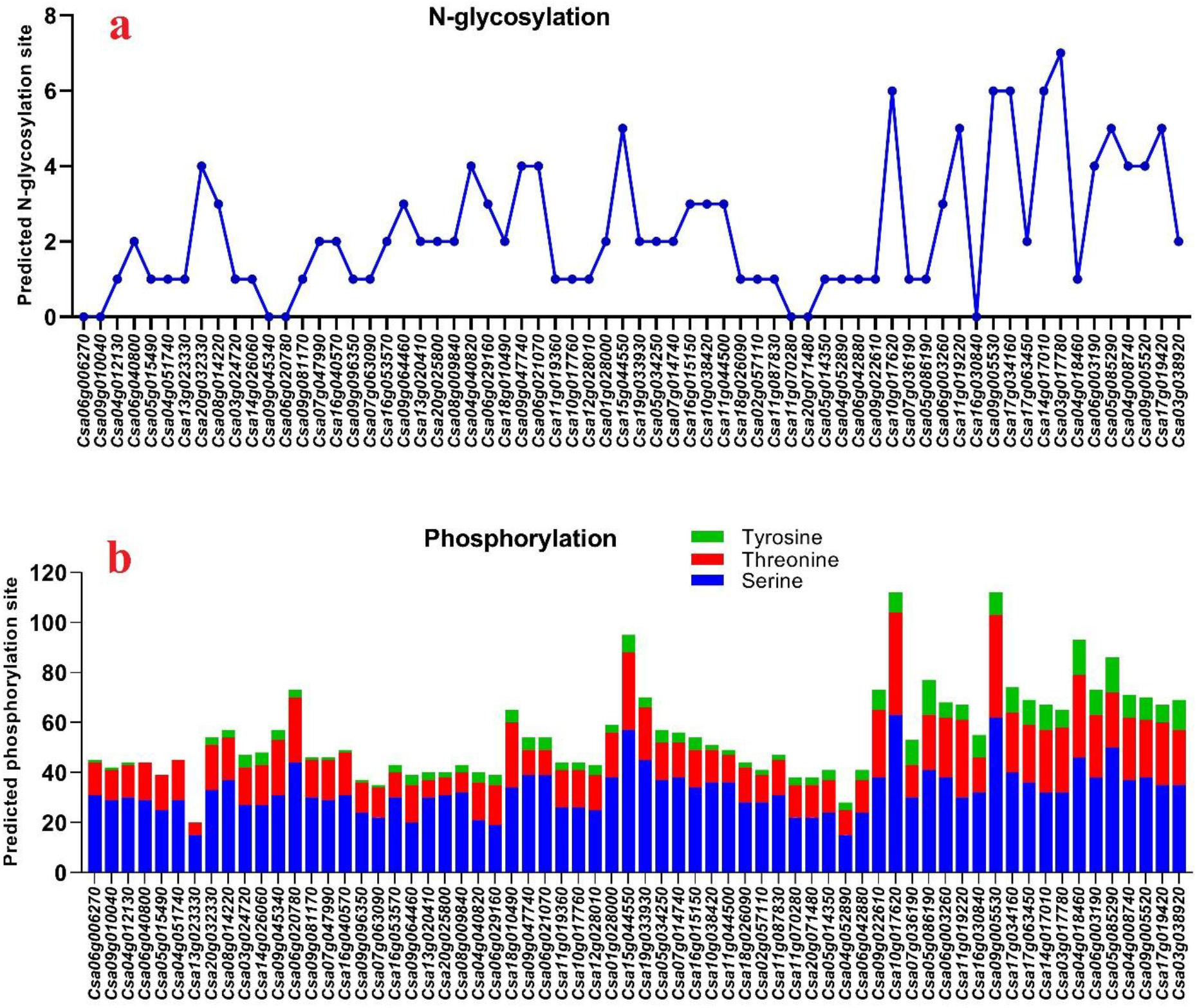
Number of predicted N-glycosylation (a), and phosphorylation (b) sites of phosphate transporter/translocator proteins in *C. sativa*

### Promoter analysis

In the present study, several key cis-acting elements related to the response of phytohormones and stress conditions were recognized in the promoter region of PHT genes (Fig. 6, 7, and Table 2). The cis-acting elements related to hormones (Table 2), including ABRE, CGTCA, TGA-element, AuxRR-core, TCA-element, P-box, and GARE-motif, were found in the promoter site of *PHT* genes (Fig. 6). The most promoter regions of the studied genes contained ABRE and CGTCA elements related to ABA and MeJA responsiveness, respectively. Besides, promoter site of four genes, including *Csa06g006270* (*PHT2;1*), *Csa18g010490* (*probable inorganic PHT*), *Csa15g044550* (*PHO1*), *Csa10g017760* (*glycerol-3-PHT*), *Csa10g038420* (*phosphoenolpyruvate translocator*), *Csa09g096350* (*glucose-6-phosphate translocator*), and *Csa05g086190* (*PHO1 homolog10*), found with high number of ABRE elements (Fig. 6). Furthermore, cis-acting elements related to the responsiveness of abiotic/biotic stresses were highly found in the promoter region of *PHT* genes (Fig. 7, Table 2). Two key cis-acting elements, MYB and MYC, were highly distributed in the upstream region of *PHT* genes. Besides, two cis-acting elements related to the responsiveness of biotic stress, including WUN-motif and W box, were identified in the promoter site of 29 *PHT* genes. Furthermore, low-temperature LTR motifs were detected in 19 *PHT* genes promoter region and drought-responsive MBS element was found in 22 *PHT* genes promoter area. In general, the results revealed that *PHTs* are involved in various processes related to growth regulation and response to adverse environmental conditions.

**Table 2.**
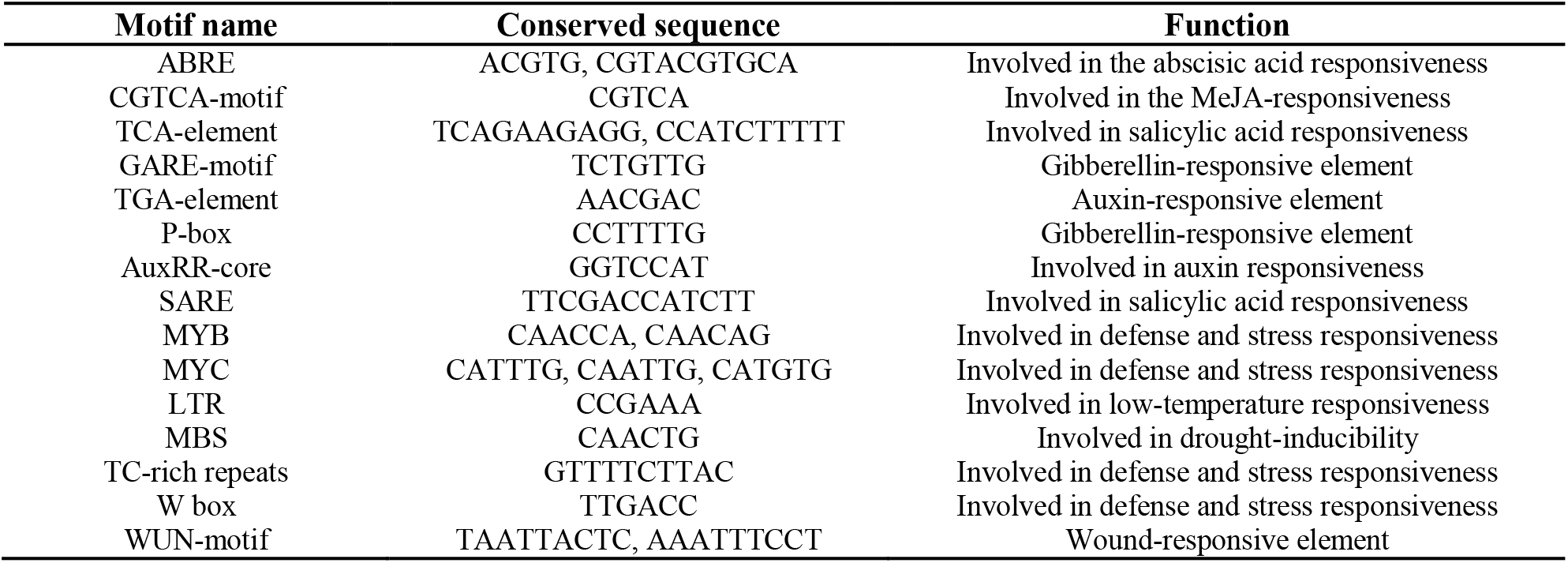
List of identified cis-acting elements in promoter site of *PHT* genes in *C. sataiva*

**Fig. 6.**
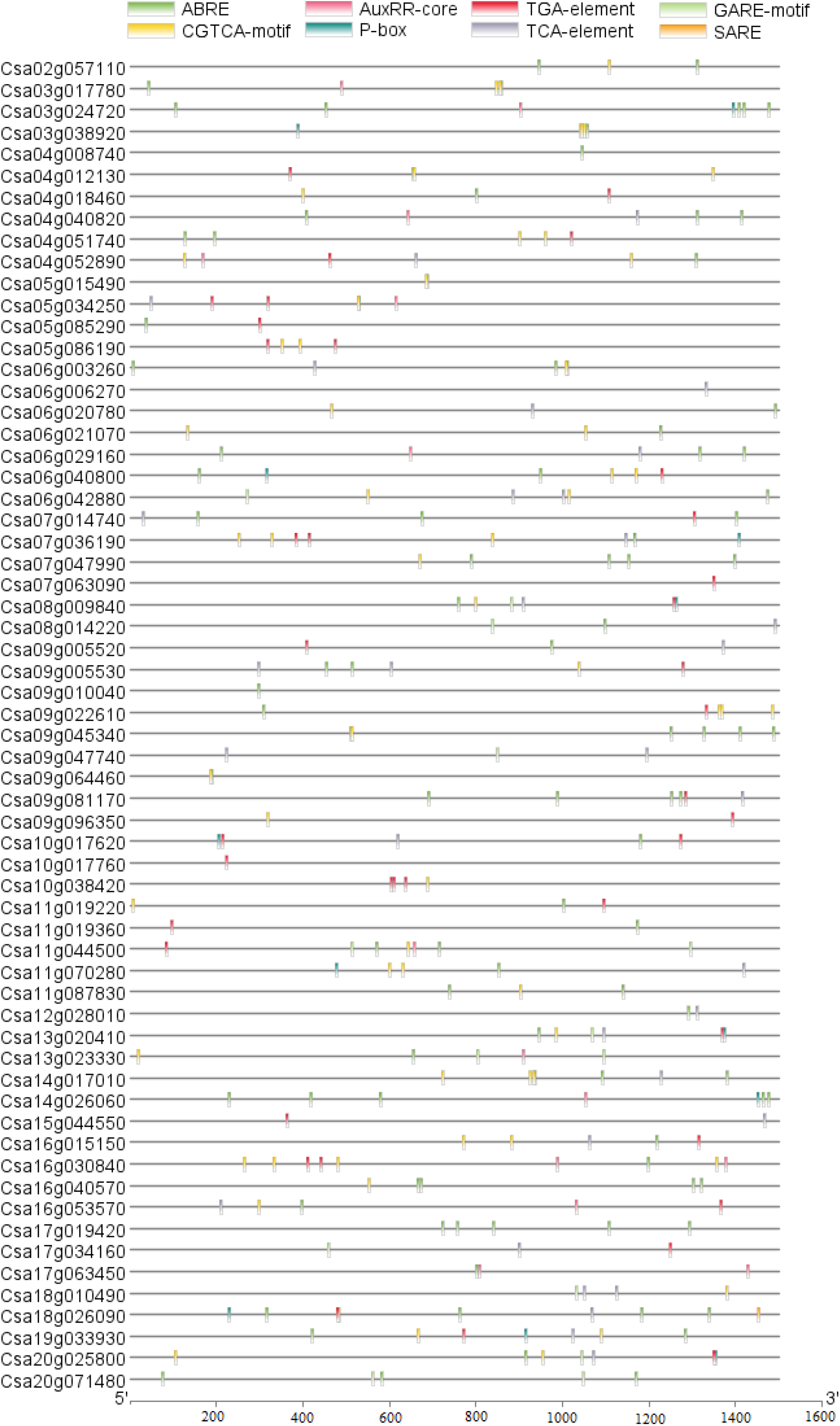
Distribution of cis-acting elements related to hormone responsive in promoter region of *PHT* genes in *C. sativa*

**Fig. 7.**
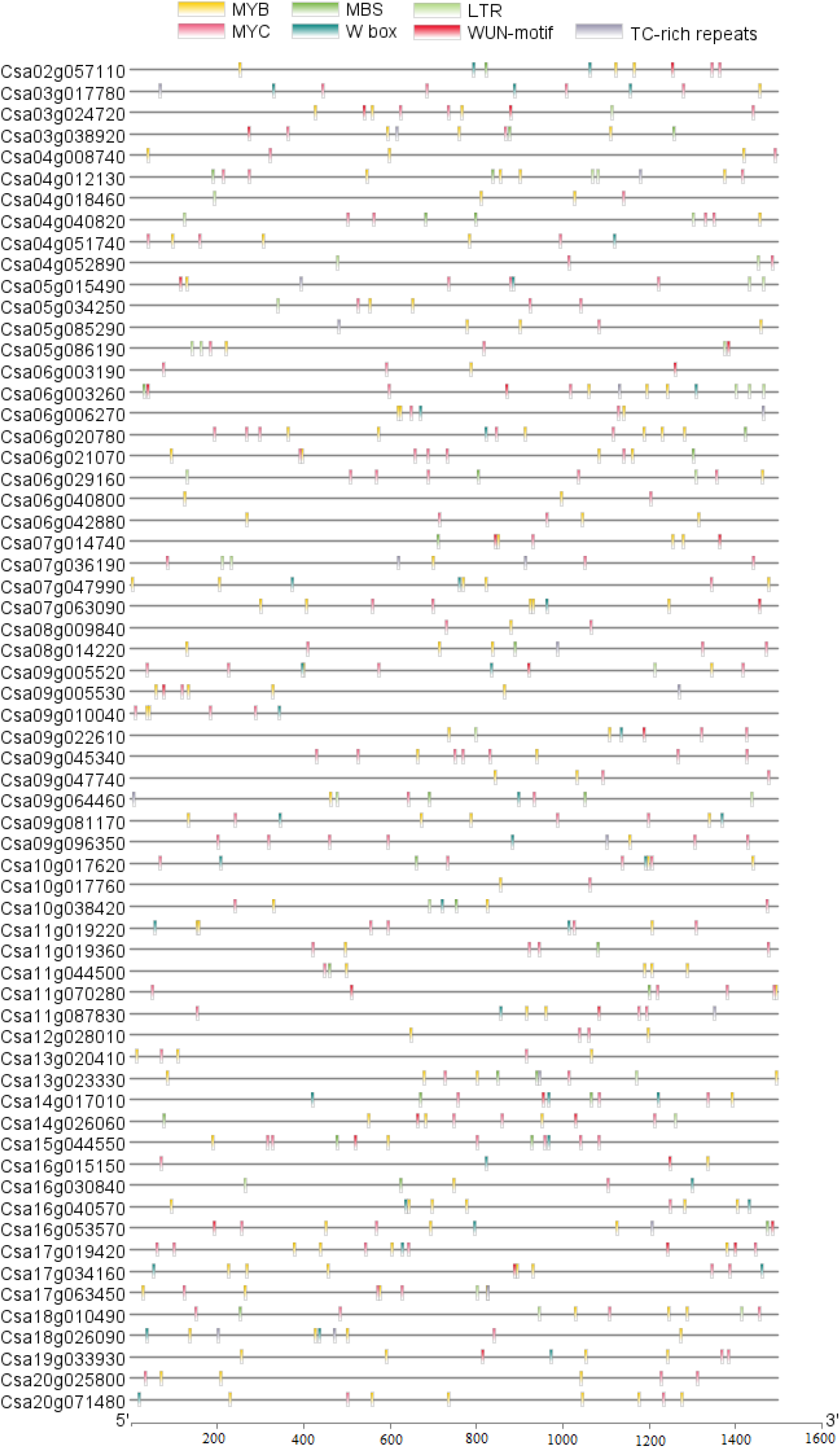
Distribution of cis-acting elements related to environmental stresses responsive in promoter region of *PHT* genes in *C. sativa*

## Discussion

Phosphorus is a key element involved in regulating growth, energy transfer, cell signaling, and increasing plant tolerance to environmental stresses (Raghothama, 2000). In plants, phosphate transporters (PHTs) are responsible for moving and distributing phosphorus in cells and organs (Smith, 2002). Several types of PHT families were recognized and characterized in different plant species, including tomato (Chen et al., 2014), rice (Liu et al., 2011), Sorghum (Wang et al., 2019), Arabidopsis (Guo et al., 2008; Wang et al., 2004), *Populus trichocarpa* (Zhang et al., 2016), *Triticum aestivum* (Teng et al., 2017), potato (Liu et al., 2017), and *Malus domestica* (Sun et al., 2017). However, in plant species such as *Camellia sativa,* the *PHT* family genes have not been widely characterized based on available genomic data. In the present study, as the first report, we found 66 *PHT* genes involved in phosphate transporter/translocate in *C. sativa*. The recognized genes were belonged to *PHT1*, *PHT2*, *PHT4*, *PHO1*, *PHO1 homologs*, *glycerol-3-PHTs*, *sodium-dependent PHTs*, *inorganic PHTs*, *xylulose 5-PHTs*, *glucose-6-phosphate translocators*, and *phosphoenolpyruvate translocators*. Regarding the physicochemical properties, the PHT proteins were stated with high diversity in terms of pI, molecular weight, GRAVY value, and exon number, indicating that these transporter genes may be associated with various cellular pathways (Ahmadizadeh et al., 2020a, 2020b). Interestingly, all PHO1 homolog proteins were showed a negative GRAVY value, revealing that the PHO1 homolog proteins are hydrophilic (Heidari et al., 2019; Rezaee et al., 2020). Besides, it was reported that PHT1 proteins are hydrophobic (Nussaume et al., 2011). More exons were observed in *PHO1* genes and their homologs that can cause different forms and functional diversity in these genes. Our findings revealed that PHTs in C. sativa with their orthologues in Arabidopsis and rice could be classified into seven different groups that PHTs in rice showed the high distance from studied dicot species. These results illustrate that probably PHT genes of dicot plants are derived from PHT genes of monocot species (Faraji et al., 2020; Heidari et al., 2020a).

The abiotic and biotic stresses reduce plant performance. Previous studies have shown that phosphate transporters are involved in response to biotic and abiotic stresses (Cubero et al., 2009; Hassler et al., 2012). In the present study, the expression analysis of *PHT* genes in various tissues of *C. sativa* revealed high expression in root tissues of *C. sativa*. These results are an agreement with previous study which also showed the high expression of *PHO1* genes in root causes phosphorus to be loaded in xylem (Hamburger et al., 2002; Liu et al., 2012; Młodzińska and Zboińska, 2016). Moreover, other *PHTs,* including *PHTs2, PHTs4,* and *PHTs3,* are more expressed in shoot tissues and involved in the distribution of phosphorus in organelles such as mitochondria, Golgi, and plastids (Gho and Jung, 2019). The RNA-seq analyses of various abiotic stresses showed role pf *PHTs* genes in drought, cold, salinity, and cadmium stress in *C. sativa*. Interestingly. However, these genes were more up-regulated in response to cold stress. It was stated that deficiency in phosphorus transfer could cause sensitivity to environmental stresses such as salinity (Cubero et al., 2009). There is a hypothesis that *PHTs* are involved in maintaining ionic balance in the cytoplasm, which probably increases the resistance of plants to stress. Under the stress of the heavy metal such as cadmium, various mechanisms have been proposed that transporters and cellular pumps play an important role in reducing the toxicity and maintaining the ionic balance (Heidari et al., 2020b; Heidari and Panico, 2020). It seems that *PHT* genes are involved in response to cadmium stress in *C. sativa*.

Regarding the Venn diagrams, we found that four *PHTs,* including a *PHT4;5* gene, a *sodium-dependent PHT* gene, and two *PHO1 homolog3* genes are up-regulated in response to all studied stresses. Interestingly, two *PHO1 homolog3* genes with high expression levels also were identified as the hyperglycosylated proteins. Glycosylation modifications as a post-translation modification can affect the stability and dynamics of target proteins Lee et al. (2015). In general, PHOs1 and their homologs were predicted with high potential N-glycosylation and phosphorylation sites. Post-translation modifications such as phosphorylation processes cause significant effects on protein activities and inducing cell signaling (Heidari et al., 2020a; Silva-Sanchez et al., 2015). The high phosphorylation potential of PHO1 proteins may indicate that these proteins are involved in various cellular processes under signal transduction pathways (Ahmadizadeh et al., 2020a). Regarding the promoter region analysis, various cis-acting elements associated with response to stress conditions and phytohormones were found in the upstream region of *PHT* genes. The cis-regulatory elements play a critical role in the regulation of gene expression that can be induced by external stimuli or induced signaling pathways in response to biotic and abiotic stresses (Ahmadizadeh and Heidari, 2014; Heidari et al., 2015; Nawaz et al., 2019). In the current study, we stated that *PHT* genes could be more induced by ABA and MeJA hormones. ABA and MeJA are two important hormones that their role is known as stress-related hormones. Besides, the promoter region of *PHT* genes contains acting-regulatory elements related to the responsiveness of low-temperature, drought and wound stress, indicating that *PHT* genes have a high potential to regulate the cellular processes associated with increasing Camelina resistance.

## Conclusion

We identified and characterized 66 *PHT* genes in *C. sativa*. Our findings revealed that *PHT* genes in *C. sativa* are diverse in functions and physicochemical properties. However, the physicochemical properties, post-translational modification, and gene expression showed that the *PHO1* genes and their homolog are more involved in different cell pathways than other types of *PHT* families. Our results may be useful for an understanding of the PHTs-mediated signaling pathways that play critical roles to increase the tolerance of plants under adverse conditions.

